# SHIP1 regulates TREM2 signalling and macrophage functions in a hiPSC-derived model

**DOI:** 10.1101/2025.07.02.662745

**Authors:** Claire S Martin, Juliane Obst, Shania Ibarra, Emma Murphy, Katerina Gospodinova, Emma Mead

**Affiliations:** ARUK Oxford Drug Discovery Institute, Nuffield Department of Medicine Research Building, University of Oxford, UK

## Abstract

Genetic and functional studies strongly implicate microglia in the pathology of Alzheimer’s disease (AD). The Triggering Receptor Expressed on Myeloid cells 2 (TREM2) pathway is an important functional regulator of microglia in AD, promoting phagocytosis of apoptotic cells, debris and pathogenic proteins including amyloid beta (Aβ). Additionally, genome-wide association studies have identified risk bearing polymorphisms in several members of the pathway including *INPP5D,* encoding the inositol phosphatase SHIP1. Recent studies utilising *in vitro* macrophage and microglia models and preclinical mouse models have identified a role for SHIP1 in both modulating TREM2 signalling and modulating neurotoxic microglial activation. In this study, we characterised the role of SHIP1 in regulating TREM2 signalling and global microglial functions using human induced pluripotent stem cell (hiPSC)-derived macrophages as a physiologically-relevant *in vitro* model of microglia. Isogenic parent and SHIP1 knockout (SHIP1 KO) lines were generated and macrophage phenotype and functions investigated. Knocking out SHIP1 was associated with a decreased expression of the lipopolysaccharide (LPS) co-receptor CD14, which translated to lower secretion of pro-inflammatory cytokines on stimulation with LPS. Additionally, SHIP1 KO hiPSC-derived macrophages showed decreased signalling upon TREM2 stimulation, which was further reflected in a reduced level of phagocytosis of apoptotic neurons compared to the parental line. The observed attenuated TREM2 signalling did not correspond to decreased TREM2 protein levels nor increased shedding of the receptor. Examination of macrophage scavenger receptor expression revealed reduced cell surface levels of CD163 and CD206 involved in resolution of inflammation but increased levels of the phagocytic receptor MerTK, while morphological analysis revealed a less amoeboid activated phenotype and increased adhesion. Taken together, our data indicates that SHIP1 plays a key role in regulating global macrophage functions in addition to modulating TREM2 signalling and inflammation.

## Introduction

Alzheimer’s disease (AD) is a progressive neurodegenerative disease and the most common cause of dementia, with an increasing incidence worldwide (1). A dysregulated neuroinflammatory response underpins AD pathology, much of which is driven by pathological activation of microglia (2). Genome-wide association studies have implicated several genes highly expressed in microglia which confer protection or risk in development of late-onset AD (LOAD). These include many components of the Triggering Receptor Expressed on Myeloid cells 2 (TREM2) signalling pathway, including TREM2 itself, Phospholipase C γ2 (PLCγ2), and Src Homology 2 (SH2) containing Inositol 5’-Phosphatase 1 (SHIP1) (3, 4).

SHIP1, encoded by *INPP5D*, is a phosphoinositide phosphatase expressed throughout cells of haematopoietic lineage, including in macrophages and microglia (5, 6). SHIP1 modulates microglial functions through regulation of phosphatidylinositol species at the plasma membrane, hydrolysing the phosphoinositide 3-kinase (PI3K) substrate phosphatidylinositol-3,4,5-triphosphate (PI(3,4,5)P)_3_ into phosphatidylinositol-3,4-biphosphate (PI(3,4)P)_2_ (7). SHIP1 thus acutely regulates the PI3K/Akt pathway and therefore cell survival, activation and metabolism (8). In addition to its function as a phosphatase, SHIP1 has been shown to regulate TREM2 signalling through binding of its SH2 domain to the immunoreceptor tyrosine-based activation motif (ITAM) of the adapter protein DAP12, preventing binding and subsequent activation of spleen tyrosine kinase (SYK) and downstream signalling (9). Genetic studies have strongly implicated SHIP1 in AD through the identification of numerous single-nucleotide polymorphism (SNP) variants. This includes the rs35349669 SNP which increases risk of LOAD, and the rs61068452 and rs10933431 SNPs which appear to confer protection (3, 4, 10, 11). Additionally, analysis of post-mortem brain samples found that *INPP5D* expression levels increased with progression of LOAD, correlating positively to amyloid plaque density (12).

Functional studies in AD mouse models have provided conflicting results. Increased *Inpp5d* expression in non-phagocytic plaque-associated microglia was reported in the 5xFAD mice (12). Additionally, loss of SHIP1 was associated with enhanced TREM2 signalling and Aβ phagocytosis, translating to reduced cognitive decline and amyloid burden in the same mouse model (13).

Meanwhile, conditional knockdown of SHIP1 in APP/PS1 mice led to increased amyloid burden despite the observed enhanced phagocytosis. This discrepancy was attributed to an increase in inflammatory microglia associated with plaques (14). *In vitro* studies in models of macrophages and microglia have sought to reconcile the conflicting outcomes of targeting SHIP1 in preclinical mouse models. Pharmacological inhibition of SHIP1 with dual SHIP1/2 inhibitors was found to promote phagocytosis of apoptotic neurons and Aβ, as well as metabolic fitness and mitochondrial activity in primary and immortalised microglial cell lines and primary murine microglia, aligning with the observed effects on amyloid burden in 5xFAD mice (15, 16). Conversely, SHIP1 agonism in BV-2 cells increased the degradation, but not the uptake of lipid-laden cargo, including apoptotic neurons and synaptosomes. SHIP1 activation was also associated with reduced inflammatory response to stimulation with both bacterial lipopolysaccharide (LPS) and Aβ (17).

The above illustrates a context-dependent role of SHIP1 in regulating microglial functioning and inflammatory profile in health and disease. To understand this better and address the conflicting results in the literature, we employed a physiologically relevant *in vitro* model of microglia: hiPSC-derived macrophages using a defined and well characterised protocol (18, 19). We performed phenotypic and functional characterisation of SHIP1 KO hiPSC macrophages to assess the role of SHIP1 in modulating TREM2 signalling and vital AD-relevant microglial functions, such as phagocytosis and cytokine release. Initial assessment revealed impaired inflammatory responses and attenuated TREM2 signalling on LPS and TREM2 stimulation, respectively. Further functional analysis showed significant reduction in TREM2-mediated phagocytosis of apoptotic SH-SY5Y neurons in the SHIP1 KO macrophages, and distinct changes in morphology and macrophage functional surface receptor levels compared to the isogenic parental control. Taken together, our results highlight an important role of SHIP1 not only as a regulator of TREM2 signalling, but of global macrophage phenotype and function.

## Methods

### Human induced pluripotent stem cell (hiPSC) lines

BIONi010-C (parent control, BioSample ID: SAMEA3158050, ECACC ID: 66540023) and BI0Ni010-C SHIP1 knock-out (SHIP1 KO) were obtained from Bioneer. The parent line BIONi010-C was derived from normal adult human dermal fibroblasts (#C-2511, Lonza) re-programmed with a non-integrating episomal vector, and is available from the European Collection of Authenticated Cell Cultures (ECACC). SHIP1 KO hiPSC lines were generated from the BIONi010-C parent line by Bioneer, using CRISPR-Cas9 mediated gene editing to target the first exon and part of the downstream intron of *INPP5D* (Suppl. Fig. 1).

The BIONi010-C SHIP1 KO clones 6 and 33 were selected for further characterisation. Cells were expanded and large-scale SNP quality-controlled batches were stored in liquid nitrogen at passage 15-25. Low passage number cells were used in all downstream experiments with a minimal number of passages post-thaw to ensure consistency.

### Culture of hiPSC-derived macrophages

hiPSCs were cultured and differentiated into tissue-resident type macrophages as previously reported (18, 19). Briefly, hiPSCs were grown on hESC-qualified Geltrex-coated cell culture plates (A1569601, Gibco) in complete mTeSR™1 media (85850, STEMCELL Technologies) and passaged as clumps using 0.5mM Ultrapure EDTA (15575020, Life Technologies) in DPBS (14190250, Gibco) a minimum of two times. Embryoid bodies (EB) were generated by seeding hiPSCs onto AggreWell-800 wells (34811, STEMCELL Technologies) in mTeSR™1 media supplemented with 50 ng/ml BMP-4 (PHC9534, Life Technologies), 50 ng/ml VEGF (PHC9391, Life Technologies) and 20 ng/ml SCF (130-096-695, Miltenyi Biotec), for 3d with daily 75% media changes. EB were transferred to low-attachment plates (657970, Greiner Bio-One) for 3d prior to transfer to T175 flasks coated with 0.1% gelatin type B (G1393-100ML, Sigma) in X-VIVO15 (LZBE02-060F, Lonza) supplemented with 2 mM Glutamax (35050038, Life Technologies), 100 U/ml penicillin and 100 μg/ml streptomycin (15140122, Sigma), 50 μM 2-mercaptoethanol (31350010, Gibco), 100 ng/ml M-CSF (PHC9501, Life Technologies) and 25 ng/ml IL-3 (PHC0031, Life Technologies). Macrophage precursors were harvested weekly and plated for 7-9 days to differentiate into hiPSC-derived macrophages in X-VIVO15 supplemented with 100 ng/ml M-CSF, 2mM Glutamax, 100 U/ml penicillin and 100 μg/ml streptomycin (macrophage media). A 50% media addition was carried out on day 3.

### Western blot

Macrophage precursors were plated at a density of 1×10^6^ cells/well in 6-well plates and differentiated for 7 days. After 7 days of differentiation, supernatant was aspirated and cells were lysed with 120 μl Pierce IP Lysis Buffer (87787, Thermo Scientific), supplemented with cOmplete protease inhibitor cocktail (4693132001, Merck) and PhosSTOP (4906845001, Merck). Lysates were centrifuged at 14,000 xg and the supernatants were collected. Protein content was quantified using Pierce Coomassie (Bradford) Protein Assay Kit (23200, Thermo Scientific), following manufacturer’s instructions.

30 μg of protein cell lysates were denatured in LDS sample buffer (NP0007, Thermo Scientific) at 70°C for 10min, then loaded into Novex 8–16% Tris–Glycine precast midi gels (WXP81620BOX, Thermo Scientific) and run at 125 V for 1.5h. Proteins were transferred to a nitrocellulose membrane using the Trans-Blot® Turbo™ RTA Mini Transfer Kit (1704271, BioRad) and the BioRad TransBlot Semi Dry System, following manufacturer’s instructions. Nonspecific binding sites were blocked with 5% BSA (05482-100G, Sigma) in TBS with 0.1% Tween20 (P9416-100ML, Sigma-Aldrich) for 1h on a shaker, then membranes were incubated with primary antibodies diluted in blocking buffer overnight at 4°C: SHIP1 (MA5-14893, T.7.7, Invitrogen, 1:1000), SHIP2 (sc-166641, Santa Cruz, 1:1000), PTEN (138G6, Cell Signaling Technology, 1:1000), TREM2 (ab209814, EPR20243, Abcam, 1:500), GAPDH (G9545, Sigma, 1:2000).

Membranes were washed 3x in TBS/0.1% Tween20 and then incubated with HRP-labelled anti-rabbit IgG (656120, Life Technologies) or anti-mouse IgG (626520, Life Technologies) at 1:5000 for 1 h at RT. Membranes were incubated with the SuperSignal West Dura Extended Duration Substrate (34075, Life technologies) and signal was detected on the FujiFilm LAS-4000 System (Raytek). Intensity of protein bands was quantified using ImageJ 1.52a software and relative quantification calculated by normalising to the intensity of GAPDH band.

### Flow cytometry for macrophage surface marker expression

Macrophage precursors were plated at a density of 1×10^6^ cells/well in 6-well plates. On day 9 of differentiation, macrophages were washed with PBS (10010015, Thermo Scientific) and detached from plates using StemPro Accutase (A6964-100ML, Sigma-Aldrich) for 10min at 37°C. Macrophages were washed with PBS, spun down at 400xg and resuspended in PBS supplemented with 1% FBS (F9665-500ML, Sigma-Aldrich) and 10 μg/ml human IgG (1-001-A, R&D Systems) to a concentration of 4×10^6^ cells/ml. Cells were incubated for 10min on ice to block Fc receptors.

Following blocking, macrophages were resuspended to 2×10^5^ cells/well in 96-well V-bottomed plates in 50 μl/well. Cells were single-stained for 30min on ice with 1 μl/2×10^5^ cells of the following PE-conjugated antibodies: CD11b (301305, clone ICRF44), CD14 (367104, clone 63D3), CD45 (368510, clone 2D1), CD163 (333605, clone GHI/61), CD204 (371903, clone 7C9C20), CD206 (321105, clone 15-2), MerTK (367607, 590H11G1E3), mouse IgG1 kappa isotype control (400114, clone MOPC-21) or mouse IgG2a kappa isotype control (clone MOPC-173; all BioLegend). Stained macrophages were washed twice with PBS and fixed with 2% PFA (sc-281692, Insight Biotechnology Ltd) for 15min at RT. Fixed cells were washed with PBS and resuspended in 250 μl/well PBS and stored at 4°C. The following day, cells were run on an LSR II with FACSDIVA software and data analysed with FlowJo (all BD Biosciences).

### Enzyme-Linked Immunosorbent Assay (ELISA)

Macrophage precursors were seeded at a density of 4×10^4^ cells/well in 96-well PhenoPlates (Revvity) and differentiated for 8 days. All conditions were run in triplicate. Macrophages were treated in with 10 ng/ml LPS (L6143-1MG, Sigma) or vehicle for 18h, with supernatants collected and stored at -20°C until analysis. For all ELISAs, plates were washed with wash buffer (PBS [P4417-100TAB, Sigma-Aldrich] with 0.05% Tween20 [P9416-100ML, Sigma]) 3x between each stage. Non-specific binding sites were blocked with blocking buffer (PBS with 1% BSA [05482-100G, Sigma]) for 1h at RT on a shaker. To develop, plates were incubated with 50 μl/well 1-Step Ultra TMB ELISA substrate solution (34028, Thermo Fisher) on a shaker. The reaction was stopped by addition of 50 μl/well 450 nm Liquid Stop Solution for TMB Microwell Substrates (LSTP-1000-01, Cambridge Bioscience Ltd,) and then absorbance read on a PHERAstar FSX (BMG Labtech). Levels of analyte produced in pg/ml were assessed based on the standard curve, with subtraction of the blank wells used to remove background signal.

For IL-6, medium-binding 96-well plates were coated with IL-6 capture antibody (14-7069-81, clone MQ2-13A5, Invitrogen) diluted to 4 μg/ml in PBS overnight at 4°C before blocking. A standard curve covering 10,000 pg/ml – 173.4 pg/ml was generated with 1.5-fold dilutions of recombinant human IL-6 (10395-HNAE, Sino Biologicals) in blocking buffer, with blank wells consisting of blocking buffer alone. LPS-treated samples were diluted 1:5 and non-LPS samples diluted 1:2 in blocking buffer. Samples and standards were incubated at RT for 2h on a shaker. Plates were then incubated with 2 μg/ml biotinylated IL-6 detection antibody (13-7068-81, clone MQ2-39C3, Invitrogen) for 1h at RT, and then 41.67 ng/ml HRP-conjugated streptavidin (N100, Thermo Fisher) for 1h at RT, washing between these stages. The plates were developed with TMB for 10min.

Levels of TNFα were measured using a human TNFα DuoSet ELISA (DY210-5, R&D Systems), following manufacturer’s instructions. Briefly, medium-binding 96-well plates were coated with 4 μg/ml TNFα capture antibody in PBS overnight at 4°C before blocking. A standard curve covering the range 1,000 pg/ml – 15.6 pg/ml was generated with 2-fold dilutions of recombinant human TNFα in blocking buffer, with blank wells consisting of blocking buffer alone. LPS-treated samples were diluted 1:40 and non-LPS samples diluted 1:2 in blocking buffer. Samples and standards were incubated at RT for 2h on a shaker. Plates were then incubated with 50 ng/ml TNFα detection antibody for 2h at RT, and then 1:40 HRP-conjugated streptavidin for 20min at RT, washing between these stages. The plates were developed with TMB for 20min.

For soluble TREM (sTREM2) levels, macrophages were incubated in fresh macrophage media for 18h to assess baseline sTREM2 release. Medium-binding plates were coated with 8 µg/ml TREM2 capture antibody (Abcam, clone EPR20243) in PBS overnight at 4°C. A standard curve covering the range 20,000 – 1,171 pg/ml was generated with 1.5-fold serial dilutions of recombinant human TREM2 (11084-H08H, Sino Biologicals) in PBS, with blank wells consisting of PBS alone. Samples were diluted 1:10 in PBS. Samples and standards were incubated on the plates for 2h at RT on a shaker. Plates were then incubated with 1.5 µg/ml biotinylated TREM2 detection antibody (BAF1828, R&D Systems) for 1h at RT, and then 41.67 ng/ml HRP-conjugated streptavidin (N100, Thermo Fisher) for 1h at RT, washing between these stages. The plates were developed with TMB for 10min.

### pSYK, pPLCγ2 and pAkt Homogeneous Time Resolved Fluorescence (HTRF)

Macrophage precursors were seeded at a density of 4×10^4^ cells/well in 96-well plates and differentiated for 8 days. All conditions were run in triplicate. HTRF was used to assess levels of phospho-SYK (Tyr525/526), phospho-PLCγ2 (Tyr1217) and phospho-Akt (Ser473) compared to total levels using Revvity kits according to manufacturer’s instructions (total SYK: 64SYKTPEG; SYK p-Y525/526: 64SYKY525PEG; total PLCγ2: 64PLCG2TPEG; PLCγ2 p-Y1217: 64PLCG2Y7PEG; total Akt: 64NKTPEG; Akt p-S473: 64AKSPEG), as previously reported (20).

Macrophages were stimulated with TREM2 activating antibody (AF18281, R&D Systems, 5 μg/ml) or IgG control (AB-108-C, R&D Systems, 5 μg/ml) for 5min at 37°C with 5% CO_2_. Treatment media was aspirated and cells were lysed in lysis buffer provided in the relevant HTRF kit supplemented with cOmplete protease inhibitor cocktail (4693132001, Merck) and PhosSTOP (4906845001, Merck) for 30min at room temperature on an orbital shaker at 220 rpm. Cell lysates were dispensed onto a ProxiPlate-384 Plus (Revvity) and incubated overnight with pre-mixed Eu^3+^-cryptate and d2 antibody diluted in detection buffer. Lysis buffer only background controls and supplemented lysate positive controls were run on each plate. Plates were read on a PHERAstar FSX (BMG Labtech) using an HTRF optic to detect emission at 665nm and 620nm. Data was exported as the RFU values of each emission and signal/noise ratio calculated in GraphPad Prism (20).

### Phagocytosis assay

Macrophage precursors were plated at a density of 3×10^4^ cells/well in 96-well PhenoPlates (Revvity) and differentiated for 8 days. All conditions were run in triplicate. SH-SY5Y cells were cultured in T75 flasks with DMEM/F12 media (11320074, Gibco) supplemented with 10% FBS (F4135, Sigma-Aldrich) and 100 U/ml penicillin and 100 μg/ml streptomycin. When confluent, SH-SY5Ys were harvested with TrypLE Express (12604013, Gibco), washed with DPBS, centrifuged at 400xg for 5 min, and re-suspended in Live Cell Imaging Solution (LCIS, A14291DJ, Invitrogen). PFA was added to a final concentration of 2%, and the cells were fixed for 10 min at RT. The cells were washed with PBS and centrifuged at 1200xg for 7 min. Fixed SH-SY5Y cells were labelled with pHrodo iFL Red STP Ester (P36011, Life Technologies, 12.5 μg per 1×10^6^ cells) for 30min at RT in low protein-binding tubes.

Labelled apoptotic SH-SY5Y cells were washed twice with DPBS, centrifuged at 1200xg for 5min and pellets frozen down at -70°C for use in phagocytosis assays within 30 days.

All phagocytosis assay treatments were carried out using X-VIVO 15 without phenol red (04-744Q, Lonza) supplemented with 2mM Glutamax and 100 U/ml penicillin and 100 μg/ml streptomycin (phagocytosis media). Macrophages were pre-treated with cytochalasin D (11330-1 mg-CAY, Cayman, 1.5 μM), an inhibitor of actin dynamics that serves as a negative control, for 1h at 37°C prior to the addition of pHrodo-labelled apoptotic SH-SY5Y cells.

pHrodo-labelled SH-SY5Y cells were thawed on ice and resuspended to 1.2×10^6^ cells/ml in phagocytosis media. Macrophage media was aspirated from all treatment wells and replaced with 100 μl/well phagocytosis media. 50 μl/well pHrodo-labelled SH-SY5Y cells were added to each well (6×10^4^ cells/well) or 50 μl/well phagocytosis media for SH-SY5Y cargo-free negative control wells. Phagocytosis was monitored using the Incucyte SX5, acclimatising the plate for 30min prior to commencing scanning. Three images per well were captured once every hour for 24h, with an acquisition time of 300ms, 10x objective, imaging both the phase and orange channels. Data was exported as macrophage confluency (phase) and total integrated intensity (TII; pHrodo fluorescence intensity at 546–568 nm excitation and 576–639 nm emission). Phagocytic capacity of macrophages was calculated in GraphPad Prism as TII/confluency and quantified as area under curve (AUC).

### Adhesion and morphology assay

Cell morphology and adhesion was assessed as previously reported (20). Macrophage precursors were plated at a density of 1×10^6^ cells/well in 6-well plates and differentiated for 8d. 96-well PhenoPlates (Revvity) were coated with recombinant human fibronectin (775306, Biolegend), truncated vitronectin (A14700, Gibco), laminin (L4544-100ull, Sigma) and collagen type I (BCO-3001-1, Enzo) at a concentration of 0.5 μg/well (1.56 μg/cm^2^) in PBS for 1h at room temperature. Coating was removed, wells washed with PBS and wells blocked with 10 mg/ml heat-inactivated BSA for 1h at room temperature. Blocking solution was removed prior to starting assay and wells washed with PBS. Control wells with neither ECM proteins nor BSA blocking and BSA blocking only were included.

Macrophages were detached from 6-well plates by incubating at 37°C with StemPro Accutase for 10min, washed and resuspended in macrophage media and then plated on coated 96-well plate at a density of 3×10^4^ cells/well. Macrophages were incubated on the plate for 3h at 37°C with 5% CO_2_. Plates were washed with PBS, fixed for 15min with 4% PFA at room temperature and then washed three times with PBS before storing in the fridge until staining.

Macrophages were permeabilised with 0.1% Triton X-100 (X100-100ML, Merck) in PBS for 15min at room temperature, washed twice and then stained for 45min at room temperature with 1:400 Alexa Fluor 488 Phalloidin (A12379, Invitrogen) and 1:2000 Hoechst33342 (H3570, Invitrogen) diluted in DPBS. Plates were washed three times with DPBS and imaged on the Opera Phenix (Revvity). 10 fields of view/well were imaged with a 40x water objective. Cell count (total number of objects from the 10 fields of view) and morphology (cell roundness and cell area in μm^2^) based on Phalloidin F-actin staining were analysed using Harmony software (Revvity), removing border objects from analysis.

### Statistical analysis

Data are shown as mean ± SEM of at least 3 independent biological repeats, and were analysed using GraphPad Prism 10 software. Normality of data was assessed using the Shapiro-Wilk normality test. Comparisons were analysed using unpaired Student’s t-test, one-way ANOVA or two-way ANOVA with multiple comparisons test as appropriate, detailed in respective figure legends. P-values from ANOVA analyses are detailed in text where appropriate.

## Results

### SHIP1 KO in hiPSC-derived macrophages leads to altered cell surface marker expression

In this study we aimed to investigate the role of SHIP1 as a negative regulator of the TREM2 pathway. SHIP1 was knocked out in the parent BIONi010-C line using CRISPR/Cas9 genome editing and two potential KO hiPSC clones were selected for initial phenotypic characterisation prior to functional analysis (Suppl. Fig. 1). Human iPSCs from parental and SHIP1 KO clones, clones 6 and 33, were differentiated to tissue resident macrophages using a well-established protocol (18, 19). Successful knockout of SHIP1 was confirmed by western blot, where both potential clones were shown to lack SHIP1 (Fig. 1 A-B). Loss of SHIP1 had no significant effect on the protein levels of related PI(3,4,5)P_3_ phosphatases, including the paralog SHIP2 (Fig. 1 C) and a more ubiquitously expressed phosphatase involved in tumour suppression and glucose metabolism, PTEN (Fig. 1 D) (21, 22).

**Figure 1:**
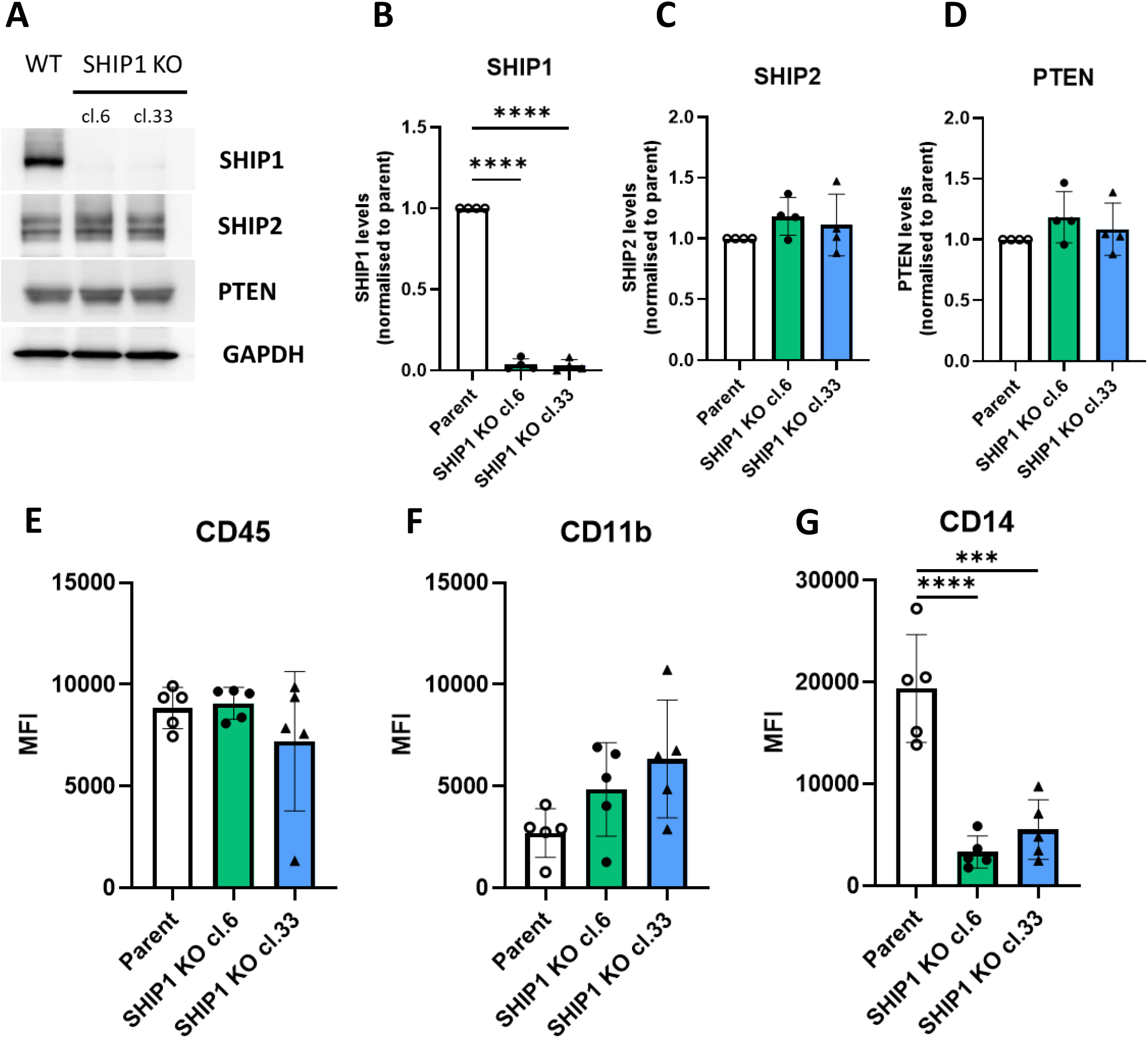
Knocking out SHIP1 in human iPSC-derived macrophages leads to altered CD14 surface marker expression. (A) Representative western blot showing expression of SHIP1, SHIP2 and PTEN in wild-type (WT) parent macrophages compared to SHIP1 KO clones 6 and 33. (B) Relative quantification of SHIP1, SHIP2 and PTEN levels in SHIP1 KO clones, normalised to that of parent macrophages. Levels of protein of interest were normalised to GAPDH expression. Surface levels of macrophage receptors (E) CD45, (F) CD14 & (G) CD11b were assessed by flow cytometry. Surface levels are represented as median fluorescence intensity (MFI). Data are N=4 (B-D) or N=5 (E-G), presented as mean ± SEM. Statistical significance of data was assessed by one-way ANOVA with Dunnett’s post-test to compare SHIP1 KO clones to parent macrophages. ***P<0.001, ****P<0.0001, ns = not significant.

The effect of SHIP1 KO on key cell surface markers for macrophages and microglia was subsequently assessed using flow cytometry (Fig. 1 E-G). Surface levels of the immune cell marker CD45 (Fig. 1 E), a protein tyrosine phosphatase, and CD11b (Fig. 1 F), an integrin involved in cell migration and adhesion, were unchanged. However, the levels of CD14 (a transmembrane co-receptor for LPS binding) were significantly decreased in the SHIP1 KO hiPSC-derived macrophages compared to the parental line (Fig. 1 G; cl.6 P<0.0001; cl.33 P<0.001). Clone 6 showed less variation in levels of CD45 and CD14, and was thus selected for further functional characterisation (CD45: coefficient of variation cl.6 8.58% vs cl.33 47.54%; CD14: coefficient of variation cl6 47.52% vs cl.33 53.27%). All subsequent reported data uses this clone unless otherwise specified.

### SHIP1 KO impairs pro-inflammatory response to LPS stimulation

The observed decrease in CD14 cell surface levels implied an altered inflammatory response to LPS stimulation in the SHIP1 KO hiPSC-derived macrophages. To investigate this further, parent and SHIP1 KO macrophages were treated with 10 ng/ml LPS for 18h and levels of the pro-inflammatory cytokines IL-6 and TNFα in the cell culture media was assessed using ELISA (Fig. 2). LPS treatment induced a significant increase in the levels of release of IL-6 and TNFα in both parent and SHIP1 KO macrophages compared to vehicle controls (Fig. 2; IL-6: P<0.001; TNFα: P<0.01). However, the effect of LPS treatment was significantly blunted in the SHIP1 KO macrophages (Fig. 2 A; P<0.01; Fig. 2 B; P<0.05), which correlates with the reduced CD14 levels observed by flow cytometry.

**Figure 2:**
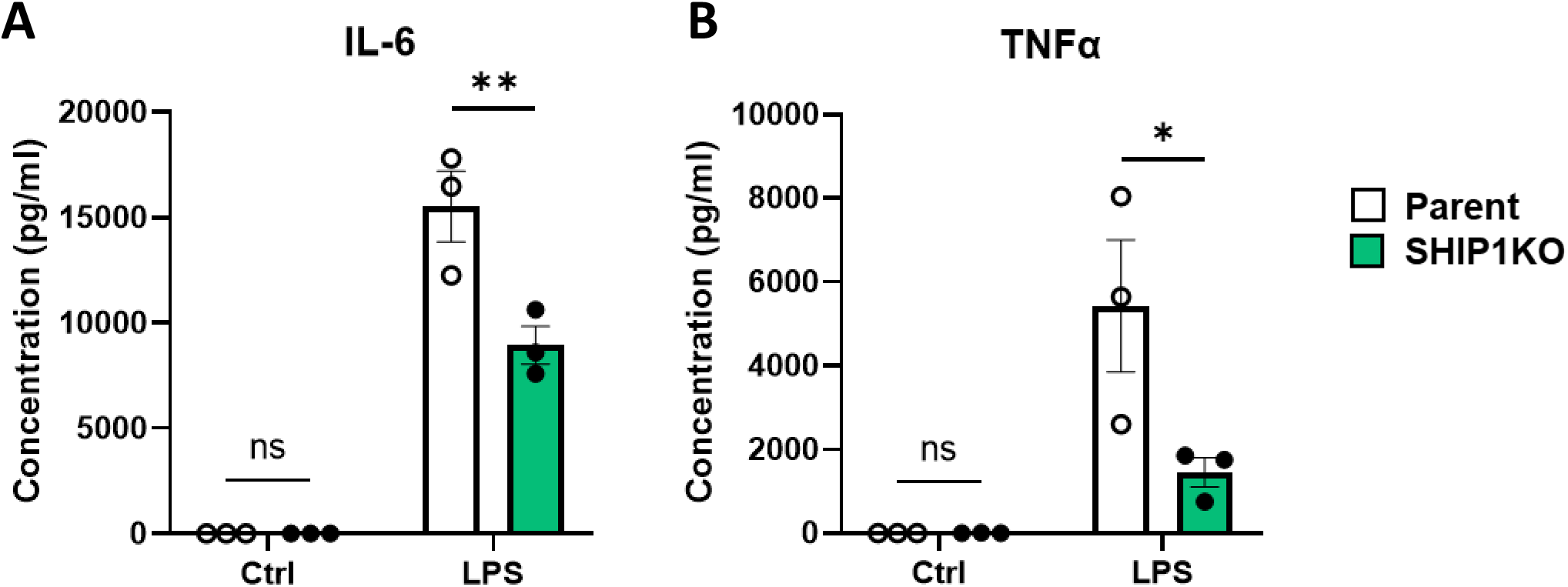
SHIP1 KO reduces inflammatory response following LPS treatment in human iPSC-derived macrophages. Parent and SHIP1 KO macrophages were treated for 18h with vehicle (ctrl) or 10 ng/ml LPS (LPS). Supernatants were collected and assessed by ELISA for levels of (A) IL-6 and (B) TNFα. Data are N=3, presented as mean ± SEM. Statistical significance of data was assessed by two-way ANOVA with Šidák’s multiple comparisons test. *P<0.05, **P<0.01, ns = not significant.

### Loss of SHIP1 attenuates TREM2 signalling in hiPSC-derived macrophages

SHIP1 has been shown to negatively regulate TREM2 signalling at multiple points: i) via competitive occupation of SYK, therefore blocking downstream phosphorylation and activation of PLCγ2; ii) through promoting hydrolysis of the PLCγ2 substrate PI(3,4,5)P_3_ into PI(3,4)P_2_ (9); and iii) through negatively regulating Akt activation (23).

We hypothesised that knocking out SHIP1 in our *in vitro* microglia-like model would lead to enhanced TREM2 signalling through increased activation of SYK, PLCγ2, and Akt. To test this, we stimulated the pathway using a TREM2 specific activating antibody (AF18281, 5 µg/ml for 5 min) and measured levels of SYK, PLCγ2, and Akt phosphorylation using HTRF. TREM2 activation was associated with a significant increase in SYK (Fig. 3 A; P<0.0001), PLCγ2 (Fig. 3 B; P<0.0001) and AKT (Fig. 3 C; P<0.0001) phosphorylation in both parent and SHIP1 KO macrophages compared to an IgG control. However, this response was significantly attenuated in the SHIP1 KO cells (Fig. 3 A; P<0.05; Fig. 3 B; P<0.0001; Fig. 3 C; P<0.001), suggestive of decreased TREM2 signalling. Baseline total levels of SYK (Suppl. Fig. 2 A; P=0.2153), PLCγ2 (Suppl. Fig. 2 B; P=0.8906) and Akt (Suppl. Fig. 2 B; P=0.6766) levels were unchanged between untreated parent and SHIP1 KO macrophages.

**Figure 3:**
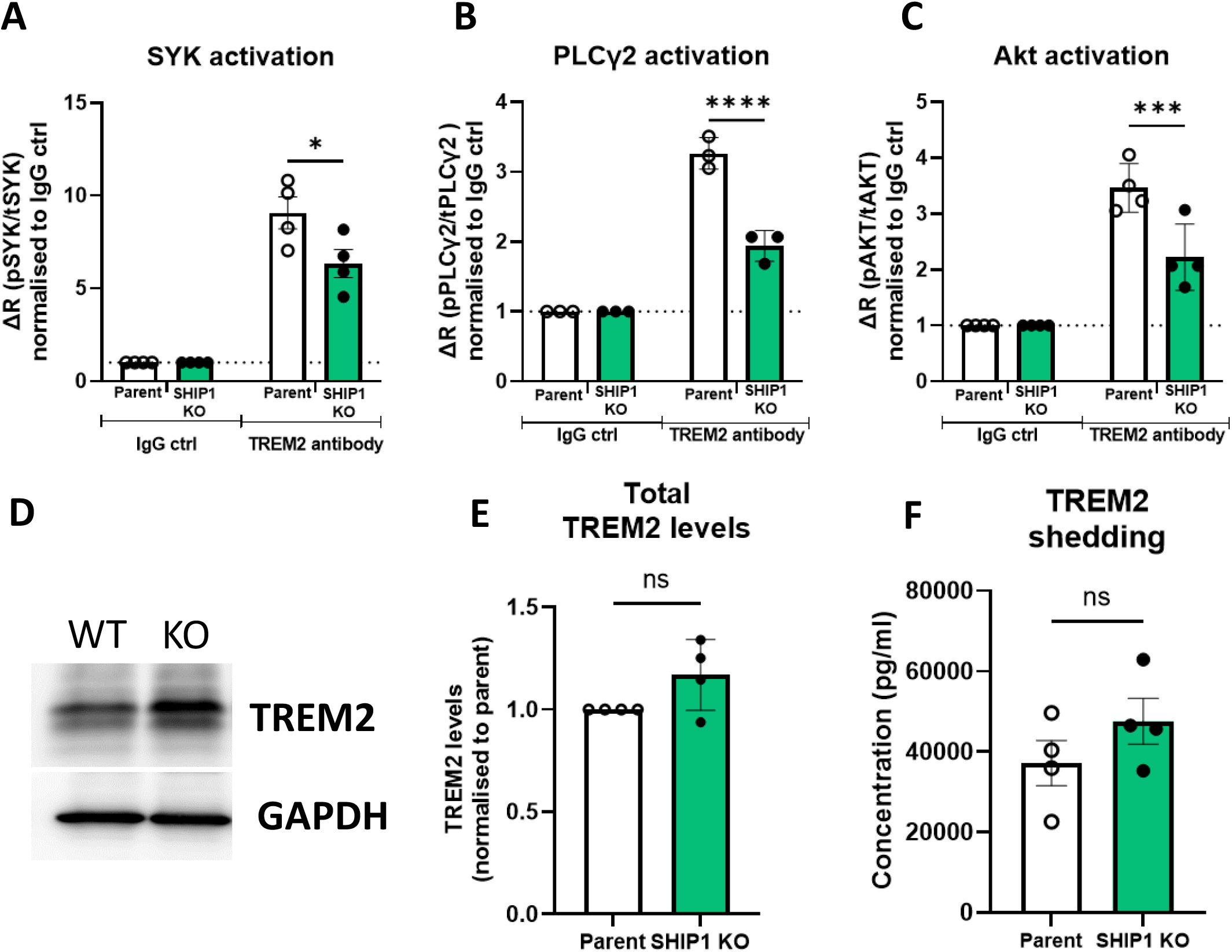
SHIP1 KO reduces TREM2 signalling in human iPSC-derived macrophages. Parent and SHIP1 KO macrophages were activated for 5min with IgG isotype control (IgG ctrl) or the TREM2-activating antibody AF18281 (TREM2 antibody), both at 5 µg/ml. The resulting phosphorylation of downstream signalling molecules (A) SYK, (B) PLCγ2 and (C) Akt was assessed by HTRF. (D) Representative western blot showing expression of TREM2 in wild-type (WT) parent macrophages compared to SHIP1 KO macrophages (KO). (E) Total TREM2 expression, normalised to GAPDH expression, in SHIP1 KO macrophages was quantified by western blot, normalised to expression in parent macrophages. (F) 18h conditioned media was collected from untreated parent and SHIP1 KO macrophages and assessed by ELISA for levels of soluble TREM2. Data are N=4 (A, C, E & F) or N=3 (B), presented as mean ± SEM. Statistical significance of data was assessed by two-way ANOVA with Šidák’s multiple comparisons test (A-C) or unpaired t-test (E & F). *P<0.05, ***P<0.001, ****P<0.0001, ns = not significant.

To investigate whether the decreased TREM2 signalling is the result of changes in TREM2 levels, we assessed TREM2 total protein levels using western blot (Fig. 3 D-E). We observed no significant changes in TREM2 protein levels upon knocking out SHIP1 (Fig. 3 E; P=0.0987). Reduced TREM2 signalling could also be the result of decreased receptor cell surface levels due to intra-membrane proteolysis (24). To assess whether SHIP1 KO alters TREM2 shedding, we assessed levels of soluble TREM2 (sTREM2) in supernatants collected from parent and SHIP1 KO macrophages by ELISA (Fig. 3 F). There were no significant differences in sTREM2 levels between the parent and SHIP1 KO macrophages (Fig. 3 F; P=0.2411).

### SHIP1 KO in hiPSC-derived macrophages reduces phagocytosis of SH-SY5Y apoptotic neurons

Phagocytic clearance of apoptotic cells is an essential function of microglia in the maintenance of brain homeostasis. We and others have previously shown that this process is largely mediated by TREM2 and requires the accumulation of PI(3,4,5)P_3_ at the phagocytic cup (25-27). Given the observed attenuated TREM2 signalling in SHIP1 KO hiPSC macrophages (Fig. 3), and the role of SHIP1 in negatively regulating PI(3,4,5)P_3_ levels (26), we investigated the ability of the SHIP1 KO hiPSC-derived macrophages to phagocytose pHrodo-labelled apoptotic SH-SY5Y neurons over 24 hours (Fig. 4). The SHIP1 KO macrophages showed a significantly reduced level of phagocytosis compared to that observed in the parent macrophages (Fig. 4 A & B, P<0.001), further confirming that SHIP1 KO confers a reduction in TREM2 signalling.

**Figure 4:**
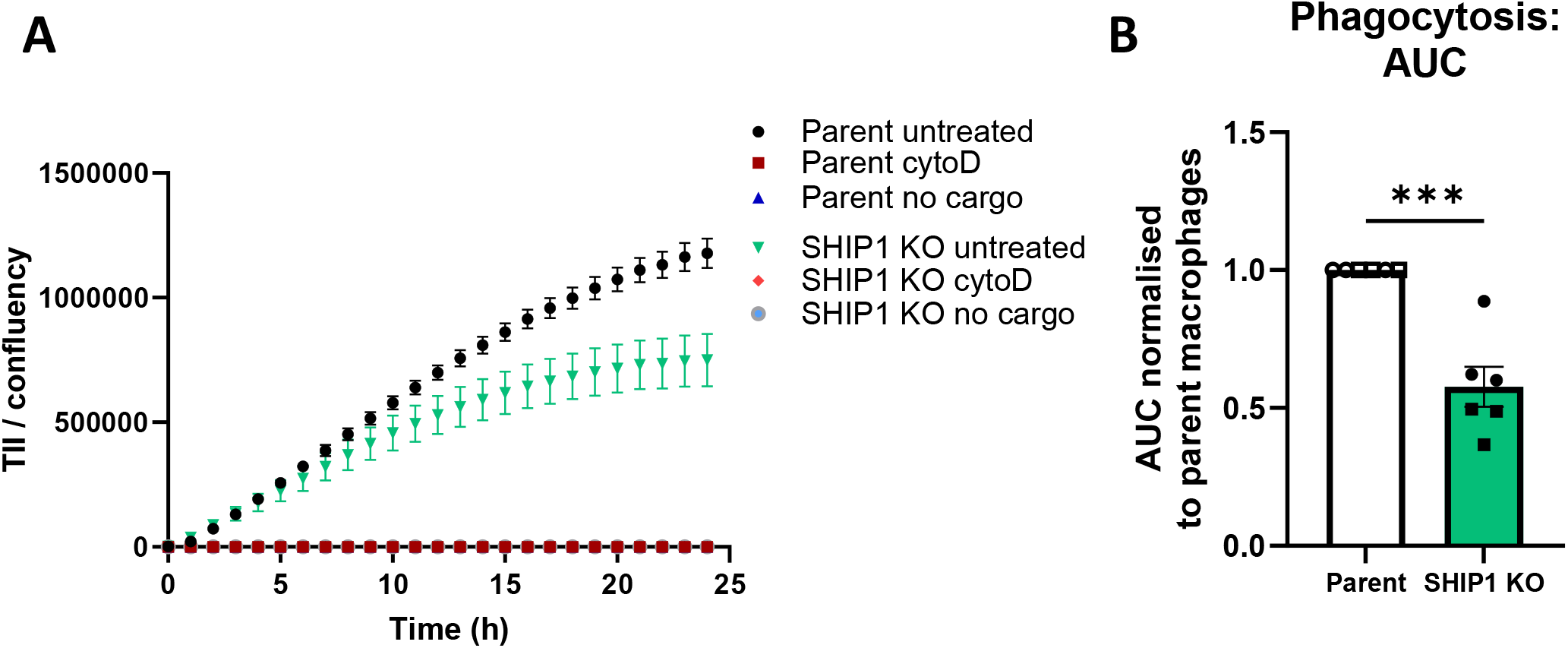
Phagocytosis is reduced in SHIP1 KO macrophages. Phagocytosis of pHrodo-labelled apoptotic SHSY-5Y neurons by macrophages was assessed on an Incucyte SX5 over 24h. (A) Representative trace of one factory set (N=3) showing levels of phagocytosis measured as total integrated intensity (TII) of pHrodo-labelled apoptotic bodies taken up by macrophages normalised to confluency of macrophage layer (TII/confluency). (B) Phagocytosis as quantified by area under curve (AUC) normalised to parent macrophages, data shows parent (white bar) and SHIP1 KO (green bar) macrophages from two independent factory sets, each of N=3 (total N=6) presented as mean ± SEM. Statistical significance of data was assessed by unpaired t-test. ***P<0.001.

### SHIP1 KO in hiPSC-derived macrophages alters global macrophage function

To further profile the effect of SHIP1 KO on macrophage phenotype and function, the expression of scavenger receptors facilitating key macrophage functions including phagocytosis and clearance of inflammatory by-products was also assessed. We examined levels of: CD163, a scavenger receptor for the haptoglobin-haemoglobin complex (28); CD206, a receptor mediating glycoprotein clearance (29); MerTK, a TAM receptor kinase regulator of inflammation and phagocytosis of apoptotic cells (30); and CD204, a scavenger receptor detecting Aβ, low-density lipoprotein (LDL), and other modified lipid proteins (31). The SHIP1 KO clones displayed decreased levels of CD163 (Fig. 5 A; P<0.05) and CD206 (Fig. 5 B; P<0.01). Conversely, the surface levels of MerTK were increased in the SHIP1 KO macrophages (Fig. 5 C; P<0.05). The expression of CD204, however, was unchanged (Fig. 5 D; P=0.6363).

**Figure 5:**
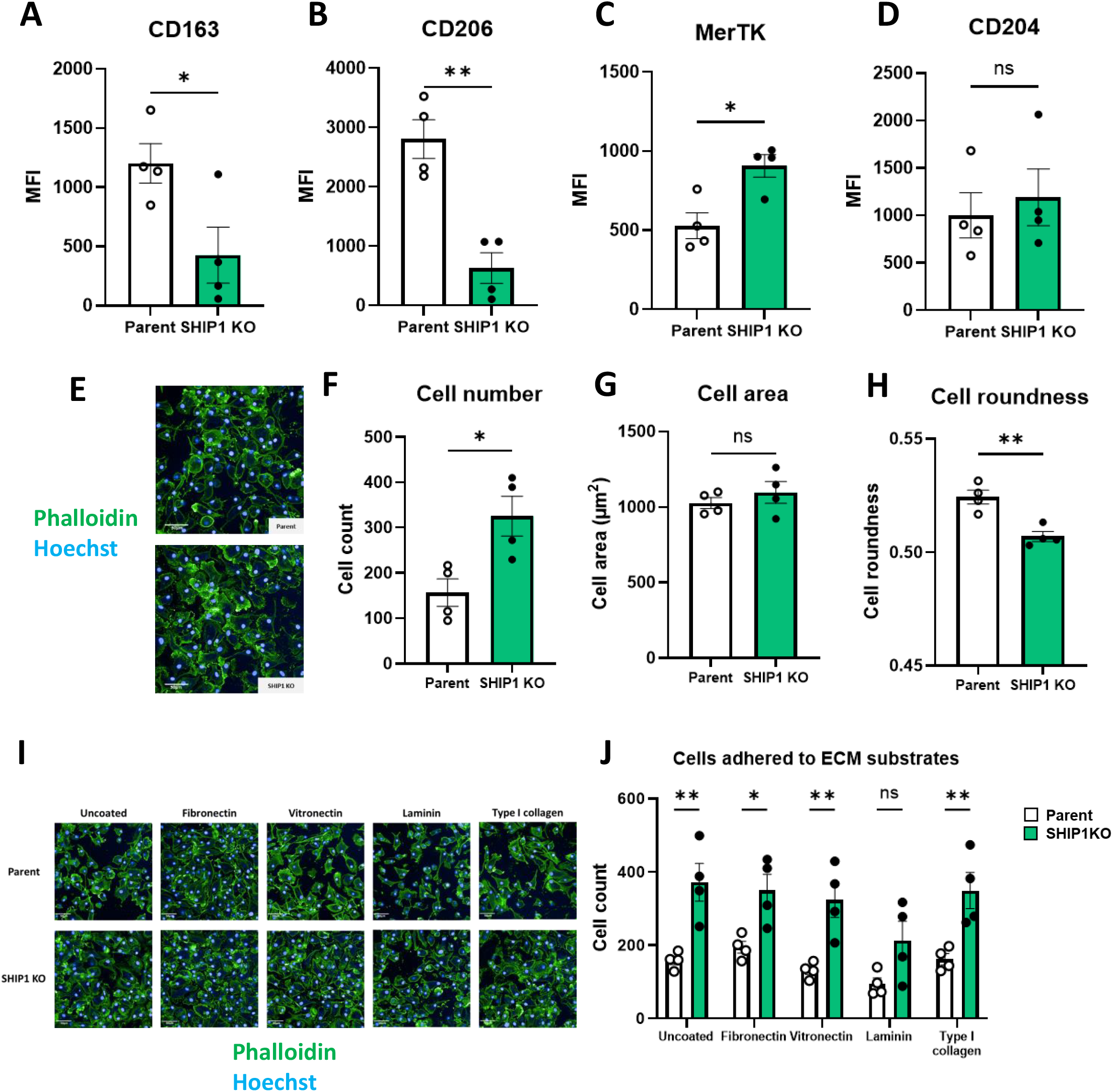
Knocking out SHIP1 in human iPSC-derived macrophages alters macrophage phenotype and morphology. Surface levels of macrophage scavenger receptors (A) CD163, (B) CD206, (C) MerTK and (D) CD204 were assessed by flow cytometry. Surface levels are represented as median fluorescence intensity (MFI). Morphology of parent and SHIP1 KO macrophages was assessed by phalloidin staining. (E) Representative image of parent and SHIP1 KO macrophages, scale bar is 50μm. Quantification of (F) cell count (number of objects in 10 fields of view), (G) cell area in μm^2^ and (H) cell roundness. (I) Representative images of parent and SHIP1 KO macrophages adhered to extracellular matrix (ECM) substrates. (J) Quantification of macrophage numbers adhered to ECM proteins. Data are N=4, presented as mean ± SEM. Statistical significance of data was assessed by unpaired t-test (A-D, F-H) or two-way ANOVA with Šidák’s multiple comparisons test (J). *P<0.05, **P<0.01, ns = not significant.

Finally, we assessed whether knocking out SHIP1 had a significant impact on macrophage morphology and adhesion in addition to the observed functional effects. Adherence and overall morphology were assessed using phalloidin staining to visualize the actin cytoskeleton. When re-plated onto untreated PhenoPlates, SHIP1 KO macrophages displayed a significant increase in cell number (Fig. 5 F, P<0.05) and reduced cell roundness (Fig. 5 H, P<0.01) compared to parent macrophages. No significant difference in cell area was observed (Fig. 5 G; P= 0.4203). To further investigate adherence, differentiated parent and SHIP1 KO macrophages were allowed to adhere to PhenoPlates left uncoated or coated with different extracellular matrix (ECM) proteins: fibronectin, vitronectin, laminin or type I collagen. Increased numbers of SHIP1 KO macrophages were found on uncoated (P<0.01) plates and on most ECM proteins (fibronectin P<0.05; vitronectin P<0.01; type I collagen P<0.01) apart from laminin (P=0.1294), indicative of enhanced adherence compared to parent macrophages.

## Discussion

Human genetics and expression data provide strong evidence for the involvement of SHIP1 in AD pathogenesis with several SNPs conferring risk, and gene expression and protein levels correlating with disease progression and pathology (3, 4, 10-12). However, downstream analysis in *in vivo* disease models have shown conflicting results, indicating that SHIP1 loss can be both neuroprotective through boosting TREM2 signalling (13), or neurotoxic by promoting inflammatory signalling (14). Both studies, however, reported enhanced microglial engagement with plaques as a result of *Inpp5d* haploinsufficiency.

Functional studies utilising *in vitro* microglial models have also attempted to determine the role and outcome of SHIP1 modulation in AD. Studies in immortalised microglial cell lines and primary murine microglia have shown neuroprotective effects of pharmacologically inhibiting SHIP1 with pan-SHIP1/2 inhibitors, enhancing TREM2 signalling and phagocytosis of dead cells and Aβ (15, 16). In contrast, activation of SHIP1 with an agonist compound in murine bone marrow derived macrophages (BMDMs) and BV2 microglial cells was shown to be beneficial by reducing the production of neurotoxic inflammatory cytokines following activation with Aβ, and promoting the degradation of phagocytosed lipid-laden cargo such as synaptosomes (17). These conflicting results may stem from the nature of the *in vitro* models used. Much of the literature describing a role for SHIP1 as a negative regulator of inflammation has been based on observations in monocyte-derived macrophage models, such as the RAW 264.7 murine cell line. These cell systems are ontogenically and transcriptionally different from brain microglia, which originate from the yolk sac, therefore could possess functional differences (16, 32-34). Other models, such as the HMC3 human microglial cell line do not endogenously express SHIP1, therefore inhibition could only be assessed following overexpression of the enzyme in these cells (16). Additionally, human and murine microglia are known to possess species-specific differences in gene expression (35), which has consequences in studies of AD-related functions and may have ramifications for SHIP1 biology. For example, human and murine microglia have been reported to respond differently to amyloid pathology: human microglia upregulate genes such as *P2RY12* and *CX3CR1* that are conversely accepted to be drivers of the homeostatic phenotype in murine microglia (36).

We sought to characterise the role of SHIP1 in TREM2-mediated signalling and downstream functions using a physiologically-relevant in vitro model of microglia: human iPSC-derived tissue resident macrophages differentiated from *Myb*-independent embryonic macrophage precursors (18, 19). We have previously used this system to study the role of PLCγ2 in TREM2 signalling and characterise the effects of missense variants implicated in AD, such as the TREM2 R47H variant (20, 27). Previous reports on the neuroprotective benefits of SHIP1 loss have identified a direct link to increased TREM2 signalling, highlighting increased levels of TREM2 and activation of SYK and Akt (13, 15). SHIP1 inhibits TREM2 signalling by binding to DAP12 and preventing PI3K and SYK recruitment and subsequent activation (23), we thus expected that genetic ablation of SHIP1 would enhance activation of TREM2-mediated downstream signalling and functions in our hiPSC-derived macrophage model. Stimulation of the TREM2 pathway using a specific TREM2 activating antibody (20), resulted in increased SYK, PLCγ2, and Akt phosphorylation compared to the corresponding IgG control in both parent and SHIP1 KO hiPSC macrophages. Unexpectedly, however, this response was significantly dampened in the KO macrophages. Analysis of TREM2 levels revealed no significant differences between the SHIP1 KO and parental macrophages, indicating that this reduction in TREM2 signalling was not a direct result of alterations in TREM2 surface levels. Nor were any changes in baseline total SYK, PLCγ2 or Akt noted between parent and SHIP1 KO macrophages, further indicating that the TREM2 pathway is preserved. The data presented here may suggest that in the absence of SHIP1 compensatory mechanisms develop to restrain excessive TREM2 signalling, particularly as this knockout was introduced at the iPSC stage prior to differentiation into macrophages. In support of this hypothesis SHIP1 is a known direct negative regulator of CSF-1R, becoming recruited to the kinase Lyn via its SH2 domain and thus antagonising Akt activation (37). Furthermore, signalling via CSF-1R synergises with TREM2 downstream of SYK, a process identified to be important for macrophage survival (38, 39). It is likely that in the context of a complete SHIP1 knockout, the hiPSC-derived macrophages adapt compensatory mechanisms to prevent excessive Akt activation as a result of altered M-CSF signalling. Future work exploring the role of SHIP1 in regulating CSF-1R signalling could help elucidate these mechanisms.

We have previously shown that phagocytosis of apoptotic neurons in our hiPSC macrophage model is predominantly TREM2-mediated, as loss of TREM2 or the downstream mediator PLCγ2 is associated with reduced levels of uptake (20, 27). In agreement with our findings of reduced TREM2 pathway activation in the SHIP1 KO hiPSC macrophages, we also observed reduced levels of phagocytosis of apoptotic neurons. SHIP1 has been consistently identified as a negative regulator of phagocytosis in different macrophage models and phagocytic contexts via the TREM2 pathway and others, including Fcγ receptors in the case of IgG-labelled cargo. This has been attributed to both SHIP1’s direct competition for binding at the ITAMs of the respective receptors as well as its metabolism of PI(3,4,5)P_3_, accumulation of which is required for phagocytic cup formation (26, 34). The only other study providing evidence for a role of SHIP1 in modulating the phagocytosis of apoptotic neurons in an hiPSC-derived microglial model has reported enhanced levels of uptake upon treatment with the pan-SHIP1/SHIP2 inhibitor, K118, but not with SHIP1 selective compounds, implicating SHIP2 in this process (15). However, in our model, knocking out SHIP1 had no effect on SHIP2 protein levels and we did not study SHIP2 in any further detail.

We observed that SHIP1 KO macrophages had increased surface levels of MerTK, a receptor highly expressed on microglia, where it binds exposed phosphatidylserine to mediate phagocytosis of apoptotic cells (30, 40), and also implicated in the uptake of Aβ and α-synuclein plaques (41, 42). MerTK has also been shown to facilitate synaptic pruning (41, 43). SHIP1 regulates receptor signalling and downstream functional effects via binding of its SH2 domain to phosphorylated ITAMs, which are not present on MerTK. The difference in MerTK levels observed in the SHIP1 KO hiPSC microglia-like model could be therefore the result of global shifts in macrophage function and/or an adaptive mechanism to compensate for the reduced TREM2 signalling. However, the increased levels of MerTK in the SHIP1 KO did not appear to be able to compensate for the loss of TREM2-mediated phagocytic capacity, at least in our assay set-up with apoptotic SH-SY5Y cells. However, we did not assess the uptake of other cargoes the uptake of which may be mediated by this differential receptor expression.

SHIP1 is well-established as a negative regulator of LPS signalling (33, 44, 45). SHIP1 can directly inhibit TLR4 signalling through blocking activation of the downstream adaptor protein MyD88 in a hypothesised competitive mechanism (33). There are no reports of a direct interaction between SHIP1 and CD14, with the latter not possessing an ITAM domain (46). However, loss of SHIP1 could potentially affect the stability of the CD14/TLR4 complex. Assessment of the inflammatory phenotype of SHIP1 KO hiPSC macrophages revealed an attenuated response to LPS, likely correlated with the reduction in CD14, a TRL4 co-receptor, in these lines. Previous work in BMDMs identified a role for SYK and PLCγ2 in mediating the endocytosis of CD14 required for TLR4 internalisation (46). It is therefore possible that the reduced levels of SYK and PLCγ2 activation we observed in the SHIP1 KO macrophages may also translate to altered CD14/TLR4 stability at the plasma membrane and thus reduced downstream signalling. Conversely, a report using *INPP5D* heterozygous loss of function hiPSC macrophages (INPP5D-HET) revealed that both acute and chronic reduction of SHIP1 activity increased NLRP3 inflammasome activation and production of the potent inflammatory cytokine IL-1β (47). 3AC treatment, however, appeared to drive a far more potent inflammatory state, including a decrease in gene expression of key scavenger receptors including CD163, CD204 and CD206. These 3AC-treated hiPSC-derived microglia also showed increased CD14 levels and spontaneous release of pro-inflammatory cytokines without inflammatory stimulation (47). In contrast, the INPP5D-HET microglia responded to the chronic loss of SHIP1 with a degree of negative feedback regulation to this inflammatory state, upregulating CD206 and the IL-1 receptor antagonist gene *IL1RN* (47). These contrasting findings are likely a result of the different models used and the more modest effect of the heterozygous knockdown compared to our own full KO. In our model of hiPSC-derived macrophages, genetic ablation of SHIP1 might promote negative feedback mechanisms to compensate for loss of associated pro-inflammatory regulation.

Similar to the effects observed upon 3AC treatment (47), we observed decreased cell surface levels of CD163 and CD206 in our SHIP1 KO hiPSC-derived microglial model. CD163 is a receptor for haptoglobin-haemoglobin, an inflammatory product of damaged erythrocytes and it has been linked to the response to small-vessel injury in AD (28, 48, 49). CD206, facilitates clearance of glycans and glycoproteins (29). Both CD163 and CD206 are associated with perivascular and meningeal macrophages. However, upregulation by microglia is commonly reported in the context of brain injury or disease (50). Downregulation upon SHIP1 KO could therefore impact the microglial response to damage and early vascular changes in dementias. These findings therefore further highlight a role for SHIP1 in regulating brain homeostasis.

Morphological analysis identified a less circular amoeboid phenotype and increased adhesion to cell culture plates, uncoated and coated with different ECM matrix proteins, in the SHIP1 KO hiPSC macrophages compared to the isogenic parent control, suggestive of a lower activation state (51). These findings are in concordance with the altered levels of scavenger receptors as well as the reduced inflammatory responsiveness we observed in the SHIP1 KO hiPSC macrophages. In our model, loss of SHIP1 was associated with a significant reduction in the cell surface levels of the LPS co-receptor CD14 and a subsequent decrease in LPS-induced inflammatory cytokine production. Despite not finding any differences in cell surface levels of CD11b, knocking out SHIP1 may lead to changes in other integrins which mediate cell attachment. SHIP1 has been reported to enhance adhesion of murine macrophages through the integrin LFA-1 (52) and inhibit chemotaxis of human blood monocytes (53). Interestingly, knockout of SHIP1 in neutrophils was found to drastically enhance their adherence through an increase in Akt activation (54). Further work exploring these effects in microglia may provide evidence for a novel role of SHIP1 in regulated microglial integrin-mediated signalling and cell adhesion. Particularly as *Inpp5d* haploinsufficiency has been shown to enhance microglial engagement with plaques, perhaps via an increase in adherence as shown in our study (13, 14).

Collectively, our data suggests an important role for SHIP1 in regulating global microglial functions in addition to modulating TREM2 signalling. Future studies should investigate the role of SHIP1 in regulating macrophage activity and function, particularly using hiPSC-derived cells that model tissue resident populations and utilising conditional knockout strategies such as CRISPR to explore effects of knockdown in differentiated populations.

## Data Availability Statement

Additional data that support the findings of this study are available online in the Supporting Information section. Data are available from the corresponding author upon request.

## Ethics Approval and Consent to Participate

The parent line BIONi010-C was generated by Bioneer from normal adult human skin fibroblasts sourced from Lonza (#CC-2511), who provide the following ethics statement: ‘These cells were isolated from donated human tissue after obtaining permission for their use in research applications by informed consent or legal authorization’.

## Acknowledgments

The authors would like to thank Mr Andrew Worth and the Jenner Institute Flow Cytometry Facility for their help with the flow cytometry work carried out in this manuscript. The authors also thank Dr Val Millar for technical assistance with imaging on the Opera Phenix.

