## Supplementary Figures for "SHIP1 regulates TREM2 signalling and macrophage functions in a hiPSC-derived model"

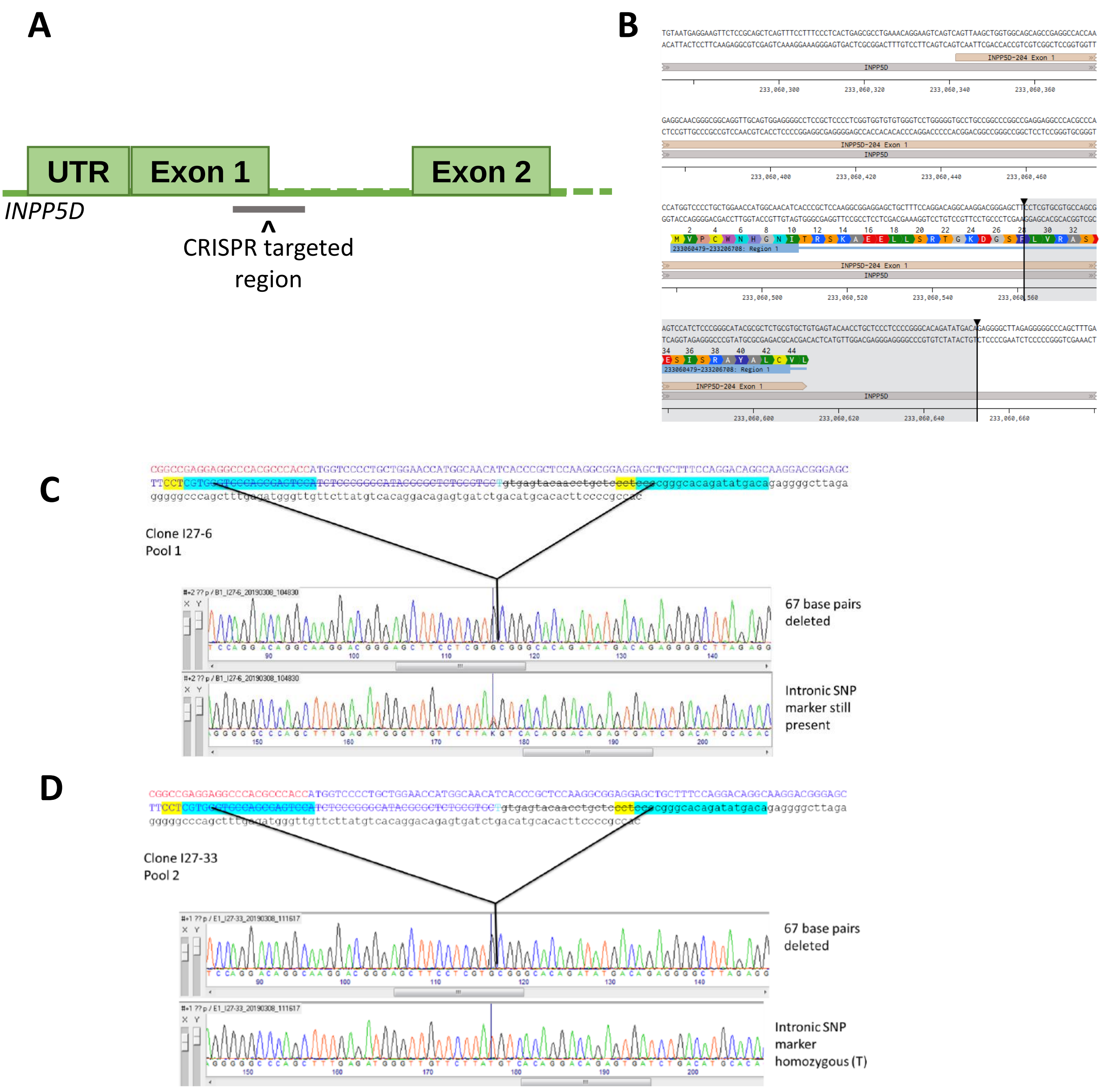


**Supplementary figure 1: Generation of SHIP1 KO iPSC lines by Bioneer**

(A) Schematic of Bioneer CRISPR knockout cloning strategy with (B) *INPP5D* gene sequence showing deleted region highlighted in grey. Sequencing analysis of SHIP1 KO clones (C) 6 and (D) 33, showing deletion of 67 bp at indicated site. The downstream intron for clone 6 shows a heterozygous SNP marker ensuring that the sequencing analysis depicts both alleles, while for clone 33 the down-stream intron shows a homozygous T at the SNP marker indicating that a larger region of the allele with the G has first been destroyed and then repaired using the allele with the T as template for homologous recombination.

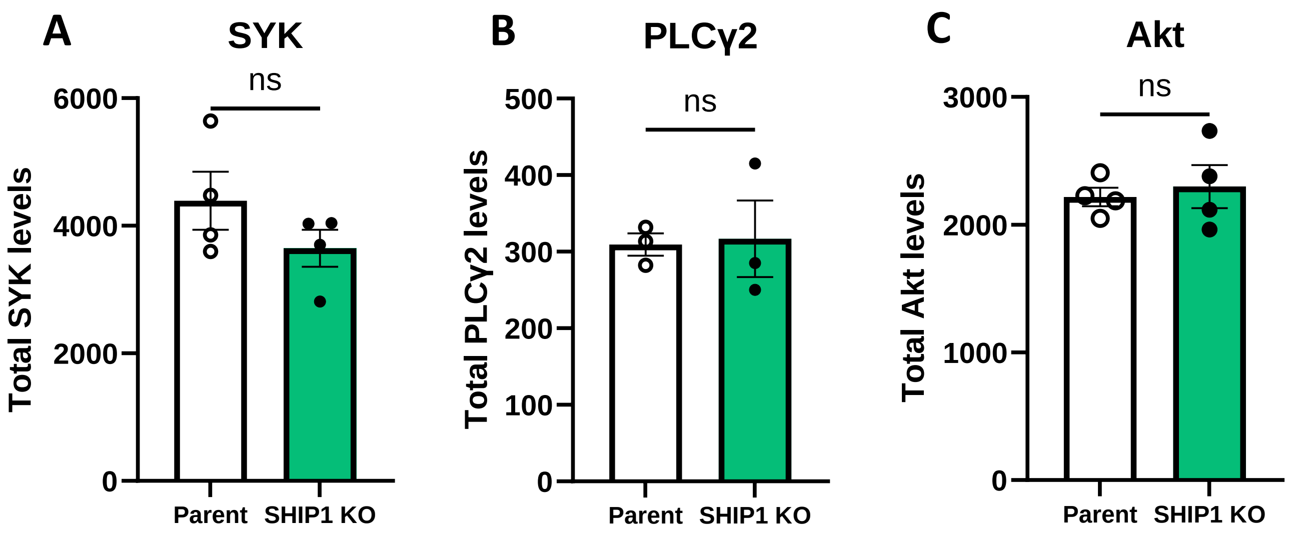


**Supplementary figure 2: Total levels of SYK, PLCγ2 and Akt**

Total baseline levels of (A) SYK, (B) PLCγ2 and (C) Akt in untreated parent and SHIP1 KO macrophages as assessed by HTRF. Data are N=4 (A & C) or N=3 (B), presented as mean ± SEM. Statistical significance of data was assessed by unpaired t-test. ns = not significant.
